# A conserved role for sleep in supporting spatial learning in *Drosophila*

**DOI:** 10.1101/2020.06.27.174656

**Authors:** Krishna Melnattur, Leonie Kirszenblat, Ellen Morgan, Valentin Militchin, Blake Sakran, Denis English, Rushi Patel, Dorothy Chan, Bruno van Swinderen, Paul J. Shaw

## Abstract

Sleep loss and aging impair hippocampus-dependent spatial learning in mammalian systems. Here we use the fly *Drosophila melanogaster* to investigate the relationship between sleep and spatial learning in healthy and impaired flies. The spatial learning assay is modeled after the Morris Water Maze. The assay uses a ‘thermal maze’ consisting of a 5×5 grid of Peltier plates maintained at 36-37°C and a visual panorama. The first trial begins when a single tile that is associated with a specific visual cue is cooled to 25°C. For subsequent trials, the cold tile is heated, the visual panorama is rotated and the flies must find the new cold-tile by remembering its association with the visual cue. Significant learning was observed with two different wild-type strains – *Cs* and 2U, validating our design. Sleep deprivation prior to training impaired spatial learning. Learning was also impaired in the classic learning mutant *rutabaga* (*rut*); enhancing sleep restored learning to *rut* mutants. Further we found that flies exhibited dramatic age-dependent cognitive decline in spatial learning starting at 20-24 days of age. These impairments could be reversed by enhancing sleep. Finally, we find that spatial learning requires dopaminergic signaling and that enhancing dopaminergic signaling in aged flies restored learning. Our results are consistent with the impairments seen in rodents and humans. These results thus demonstrate a critical conserved role for sleep in supporting spatial learning, and suggest potential avenues for therapeutic intervention during aging.

**STATEMENT OF SIGNIFICANCE:** We have studied the relationship between sleep and plasticity using a Drosophila learning assay modified after the Morris Water Maze. Using this assay, we find that sleep loss impairs spatial learning. As in mammals, flies exhibited age-dependent spatial learning impairments. Importantly, the age-dependent impairments were reversed by enhancing sleep. Interestingly, our results mirror studies on hippocampus dependent memories in rodents and humans. Thus, our data describe an evolutionarily conserved role for sleep in regulating spatial learning. They also support augmenting sleep as a therapeutic strategy to ameliorate learning impairments.

## INTRODUCTION

While a precise function of sleep remains unclear ^1^, many lines of evidence point to a pivotal role for sleep in supporting learning and memory ^2-4^. Further, cognitive impairments associated with aging and neurodegenerative disorders are associated with defects in sleep ^5, 6^. Understanding how sleep benefits neural function thus has the potential to not only reveal novel insights into brain function, but also to suggest avenues for therapeutic intervention in animals whose nervous systems are challenged by aging or neurodegenerative diseases.

In humans, sleep supports many kinds of memories ^7-10^. However, declarative memories – memories of experiences (episodic memory) and memories of facts (semantic memory), appear to particularly benefit from sleep ^11^. Importantly, sleep supports both the encoding of new information ^12, 13^, and the consolidation of learned information into a memory^11^. Further, defects in encoding new declarative memories such as new facts or names are a common feature of cognitive decline in aging and degenerative disease ^5, 14^.

Studying declarative memories in animal models remains challenging. However, rodent spatial learning and human declarative memories share common cellular substrates and computations leading to the proposal that rodent spatial learning is an evolutionary precursor of human episodic memory ^15, 16^. A spatial learning assay has been described in flies ^17, 18^. Here we have adapted this spatial learning assay for sleep-plasticity studies, and use it to investigate the effects of enhancing sleep on learning impairments resulting from aging and the classic memory mutant *rutabaga* (*rut*).

## Methods

### Flies

Flies were cultured at 25°C at ∼50% relative humidity, and reared on a standard yeast, corn syrup, molasses, and agar diet while being maintained on a 12hr light: 12 dark cycle. Female flies were used as subjects in most experiments except for the experiment with *rut*^2080^ flies in figure 2, where male flies were used.

**Figure 1.**
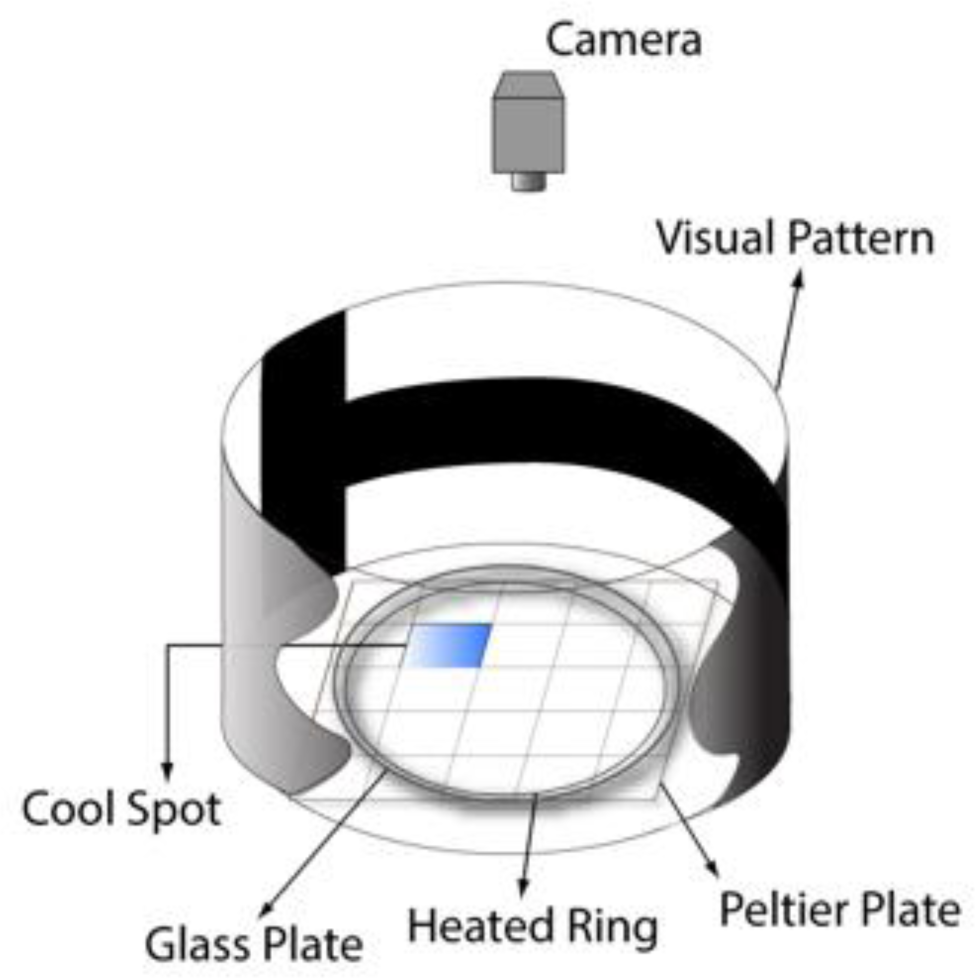
Spatial Learning apparatus. The floor of the apparatus is made up of a 5×5 grid of Peltier plates, which are maintained at a temperature of 36-37°C (which is aversive to flies). The first trial begins when a single fly in placed into the apparatus and one of the tiles is cooled to 25°C via an Arduino Uno controller (not shown). Distal visual cues mark the cool spot. In subsequent trials the location of the cool spot and the distal visual cues move in tandem such that the fly learns to associate the visual cue with the location of the cool spot. The location of the flies is monitored using a camera (see methods for details).

**Figure 2.**
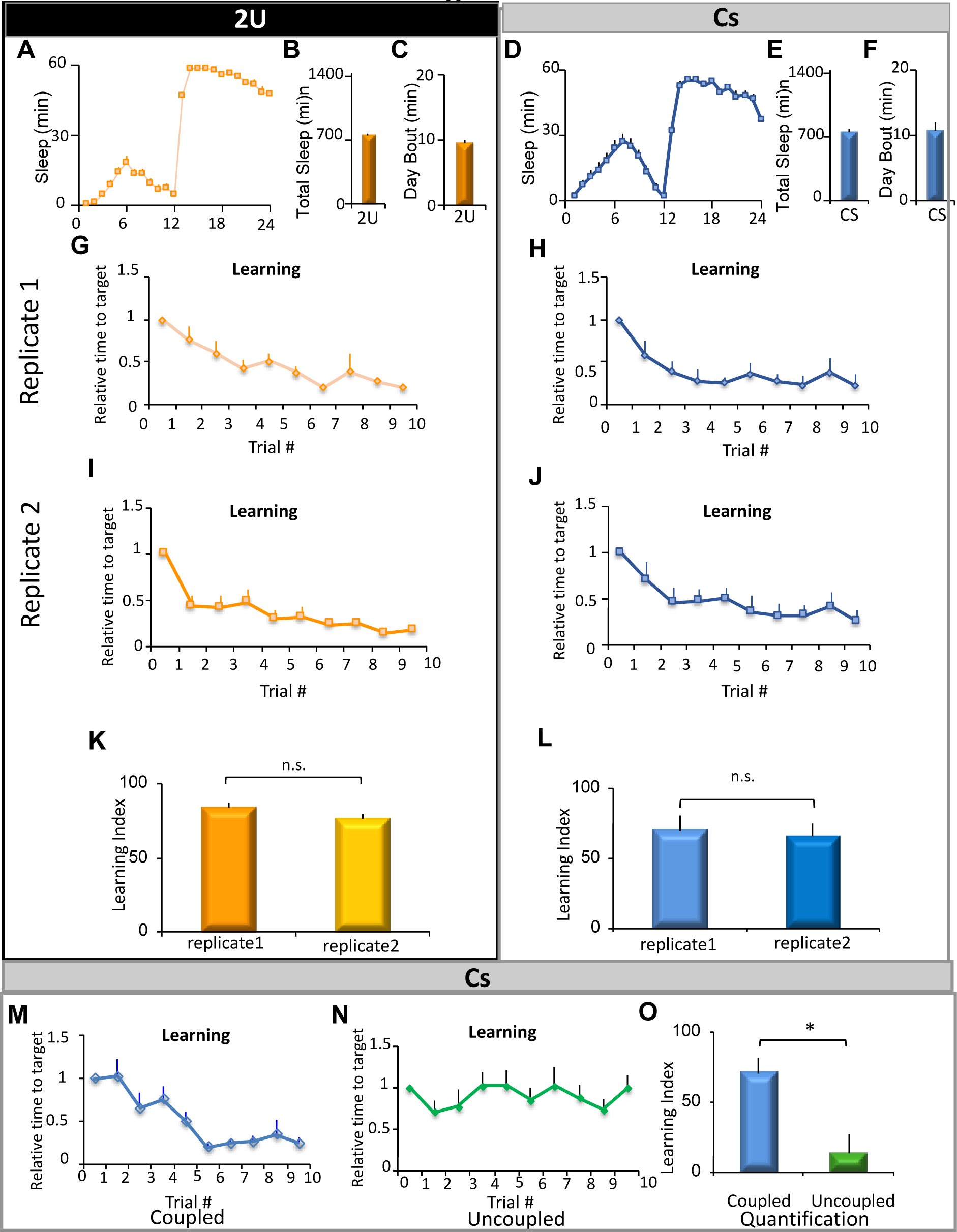
Validation of our spatial learning apparatus. (A) Sleep in minutes per hour for the 2U wild-type strain maintained on a 12:12 Light Dark schedule (LD) (*n* = 29 flies) (B) Total sleep time in minutes in 2U flies (C) Average daytime sleep bout duration (a measure of sleep consolidation during the day) in 2U flies. (D) Sleep in minutes per hour for the *Cs* wild-type strain maintained on a 12:12 LD schedule *(n* = 32 flies) (E) Total sleep time in minutes in *Cs* flies (F) Average daytime sleep bout duration in *Cs* flies. (G and I) Spatial learning in two independent cohorts of 2U flies trained in the coupled condition. Spatial Learning is expressed as the ‘time to target’ normalized to the time in the first trial. Flies reduced their ‘time to target’ over 10 trials by ∼80% (*n* = 9-11 flies/replicate, Repeated Measures ANOVA for Trials, F_[9,162]_=15.36, *p* < 10^−10^). (K) Quantification of learning scores in G & I, expressed as percentage change in the time to target in trials 9 & 10, relative to trial 1. The two replicates of 2U flies exhibited similar Learning Indices (n.s. *p* = 0.09, 2 tail t-test) (H and J) Learning in two independent replicates of *Cs* flies trained in the coupled condition. Flies reduced their ‘time to target’ over 10 trials by ∼70% (*n* = 8-10 flies / replicate, Repeated Measures ANOVA for Trials F_[9,160]_=, p < 10^−8^). (L) Two independent cohorts of *Cs* flies exhibited similar Learning Indices (n.s. *p* = 0.84, 2 tail t-test) (M and N) In contrast to flies trained in the coupled condition, *Cs* flies in the uncoupled condition showed little to no improvement in their time to target (*n* = 7-12 flies / condition, Two way repeated measures ANOVA for Condition X Trial F _[9,198]_=4.93, p= < 0.01) (O) Learning Index of flies in the ‘coupled’ condition is much higher than in the ‘uncoupled’ condition (* *p* < 0.01, t-test).

### Fly strains

*Cs* flies were obtained from T. Zars (Univ. of Missouri). 2U and *rut*^2080^/*FM7c* flies were gifts of J. Dubnau (Stonybrook University, NY). *UAS Kir2*.*1EGFP* (homozygous viable 3^rd^ chromosome insert) was a gift of R. Baines (Manchester). *TH* GAL43 (3^rd^ chromosome insert) *R15B07* GAL4, *tubPGAL80*^ts^ (3^rd^ chromosome insert) and *UAS NaChBacEGFP 4* (homozygous viable 2^nd^ chromosome insert) flies were obtained from the Bloomington *Drosophila* Stock Center. *dumb*^2^ (*Dop1R1*^f02676^) flies were obtained the Exelexis collection.

### Drug Feeding

Gaboxadol (Sigma-Aldrich, St Louis, MO) was fed to flies at a concentration of 0.1 mg /ml dissolved in standard fly food as previously described ^19^. 3IY (Sigma-Aldrich, St Louis, MO) was administered in the food at 10 mg/ml, and L-Dopa (Sigma-Aldrich, St Louis, MO) was dissolved in the food at 5 mg / ml as per established protocols ^20^.

### Sleep

Sleep was measured using protocols previously described ^21^. Briefly, individual flies were aspirated into 65mm glass tubes with standard fly food at one end, and their locomotor activity was continuously monitored using the *Drosophila* Activity Monitoring (DAM) System (Trikinetics, Waltham, MA). Locomotor activity was binned in 1min intervals; sleep, defined as 5min of inactivity, was computed using custom Excel scripts. In sleep plots, sleep in min/hour is plotted as a function of Zeitgeber Time (ZT). ZT0 represents the beginning of the fly’s subjective day (lights on), and ZT12 represents the transition from lights on to lights off.

### Sleep Homeostasis

4-7 day old female flies were placed in DAM tubes and their sleep was recorded for 2 days to establish a baseline. Flies were then sleep deprived for 12 hours during the dark phase (ZT12-ZT0) by placing DAM monitors in the Sleep Nullifying APparatus using procedures previously described ^20^. For each individual fly, the difference in sleep time in the two recovery days and the baseline day was calculated as the sleep gained / lost.

### Visual Learning Protocol

We constructed a visual place learning assay modeled on the classic Morris water maze (Figure 1) ^17, 18^. Our assay uses a ‘thermal maze’ consisting of a grid of Peltier plates maintained at 36-37°C (which is aversive to flies), and a distal visual panorama. One of four tiles can be cooled to ∼25°C. The visual panorama was arranged such that the edge between the horizontal bar panel and the vertical bar panel marked the cool spot. The experimental protocol consisted of ten 3-min training trials with a 1 min break between trials. Software written in Processing generated a random list of 10 cool spot locations.

The first trial begins when a single tile that is associated with the visual cue is cooled to 25°C. For subsequent trials, in the coupled condition, the previously cold tile is heated, and the visual panorama is rotated such that the flies must find the new cold-tile by remembering its association with the visual cue. In the uncoupled condition, cool spot locations were changed between trials but the distal visual cues remained fixed to the location in the first trial. To evaluate learning, an individual fly is placed into the apparatus and the time to find the cool spot is calculated. Individual flies remained in the arena for the duration of the experiment (10 trials). Experimenters were blinded to condition / genotype. Learning during subsequent trials, expressed as the time to target, was normalized to the time to find the cold spot in the first trial. Further, a learning index was computed as the percentage change in relative time to target as: Learning Index = (1-[relative time to target in trials 9 & 10])*100.

### Construction of Arena

The design of our visual place learning apparatus and visual panorama was adapted from previously described designs ^17, 18^. The floor of the apparatus constituted a thermal maze and was composed of twenty-five 40mmX40mm Peltier devices (Custom Thermoelectric # 12711-5L31-06CQ, Bishopsville, MD) arranged in a 5×5 grid. This Peltier grid was covered by white masking tape to create a uniform surface. The grid was connected in five groups which were soldered in series in each group to reduce the difference in temperature between the first and the last components. Four Peltier tiles of the central 9 tiles can change their state independently from cooling to heating by relays. The temperature is measured by a thermocouple and sampled with an Arduino Uno. The micro-controller controls the temperature with a power supply by changing the constant current going through the Peltier elements. The Arduino Uno also controls which Peltier device to change from a heating to cooling state. The Peltier array was maintained at 36°C-37°C except for the four tiles which could be selectively cooled to ∼25°C.

Flies were confined to this arena by means of a heated 3mm high, 200mm diameter aluminum ring that circumscribed the arena. The ring was connected by means of insulated wire to a power supply (BK Precision 1685B), which ensured that the ring was heated to 50°C, thus keeping the flies away from the walls. A glass dish coated with the siliconizing reagent Sigmacote (Sigma-Aldrich, St Louis, MO) was placed on top of the ring. The distal visual cues used for place learning consisted of one panel of alternating black & white vertical bars, one panel of alternating black & white horizontal bars, and one panel of alternating black & white angled bars printed on white paper and held together with clips. When viewed from the arena’s center the width of each bar spanned 15°. The arena was illuminated with white light, and the fly’s position was recorded with a webcam (Logitech 270).

### Heat Avoidance

Flies were confined to a chamber spanning the dimensions of 2 Peltier tiles using Lego bricks (Billund, Denmark). One of the tiles was maintained at 36-37°C, and the other at 25°C. The walls of the chamber were coated with Sigmacote (Sigma-Aldrich, St Louis, MO) to prevent flies from climbing on the sides. Flies thus had to choose between the hot side and the cool side. Heat avoidance index was calculated as the fraction of time flies spent on the cool side in a 3 min trial. Wild-type flies typically spent ∼80-90% of the time on the cool side. As a control we also tested flies when both Peltier tiles were at the same (hot) temperature. In this condition, flies did not display a preference for either side (data not shown).

### Optomotor

Flies had their wings clipped on CO_2_, at least 2 days prior to the experiment. During the experiment, flies walked freely on a round platform, 86mm in diameter, surrounded by a water-filled moat to prevent escape. Experiments using moving gratings were conducted with clockwise and anticlockwise gratings for 1.5min each. Independent flies were used for each 3-minute experiment. The temperature of the arena was 24-26 °C during experiments. The walls of the arena consisted of 6 LED panels that formed a hexagon surrounding the moat (29cm diameter, 16cm height), and onto which the visual stimuli were presented. Each LED panel comprised 1024 individual LED units (32 rows by 32 columns) and was computer-controlled with LED Studio software (Shenzen Sinorad, Medical Electronics, Shenzen, China). A camera (SONY Hi Resolution Colour Video Camera CCD-IRIS SSC-374) placed above the arena was used to detect the fly’s movement on the platform at 30 frames per second, and open-source tracking software was used to record the position of the fly (Colomb et al, 2012). All visual stimuli were created in VisionEgg software (Straw, 2008), written in Python programming language. The refresh rate was 200hz. The luminance of the LED panels was approximately 770 Lux, reaching 550 Lux at the centre of the arena. A grating of alternating cyan and black stripes were rotated in either direction (1.5min each), with a temporal frequency of 3hz and spatial frequency 0.083 cycles/degree. Analyses were performed using CeTran (3.4) software (Colomb et al, 2012), as well as custom made scripts in R programming language. For optomotor responses, the angular velocity (turning angle/second) in the direction of the moving grating was calculated.

### Statistical Analysis

Data are presented as the average accompanied by the SEM (Standard Error of the Mean). Statistical analyses were carried out in Systat software. Statistical comparisons were done with a Student’s t-test or, where appropriate, ANOVA followed by modified Bonferroni test comparisons; significance was defined as p<0.05.

## RESULTS

To expand the tools available to elucidate the molecular mechanisms underlying sleep and plasticity, we created a modified version of a *Drosophila* visual learning assay that has similarities to the Morris water maze ^17, 18^. The assay uses a ‘thermal maze’ consisting of a 5×5 grid of Peltier plates maintained at 36-37°C (which is aversive to flies) (Figure 1). The first trial begins when a single tile that is associated with a specific visual cue is cooled to 25°C. For subsequent trials, the cold tile is heated, the visual panorama is rotated and the flies must find the new cold-tile by remembering its association with the visual cue. Over the course of ten trials, flies get progressively faster at locating this cool spot ^17^.

To validate our modified Spatial Learning apparatus, we evaluated behavior in Canton-S (*Cs*) and 2U flies. *Cs* and 2U flies are frequently used as wild-type strains in sleep and memory studies, respectively ^22-24^. As seen in Figure 2 A-F, sleep characteristics of 2U and *Cs* flies were in the range observed for wild-type flies ^25^ (Figure 2, A-F). To evaluate learning, an individual fly is placed into the apparatus and the time to find the cool spot is was calculated. Learning during subsequent trials, expressed as the time to target, is normalized to the time to find the cold spot in the first trial. As seen in Figure 2G, 2U flies reduced their time to target by ∼80% over 10 trails, consistent with previous observations ^17^. To evaluate the robustness of this assay in our lab, we evaluated learning in an independent cohort of 2U flies and found similar results (Figure 2I). To simplify comparisons, we calculate a learning index (1-[average time to target in trials 9 & 10])*100. As seen in Figure 2K, the two independent replicates of learning in 2U flies were not statistically different even though the experiments were conducted on separate cohorts of flies evaluated weeks apart. Importantly, *Cs* flies showed similar learning profiles and this pattern of behavior was also observed in an independent cohort (Figure 2 H,J,L). Thus, both 2U and *Cs* flies get progressively faster at locating the cool spot.

Although these data are consistent with previous results and suggest that flies are learning the location of the cool tile in relation to a visual cue, it is possible that the flies are using other cues to improve the speed with which they can escape the heated tiles that are not dependent on learning the association between the visual panorama and the location of the cold tile (e.g. self-motion cues, undetectable thermal gradients, etc.). To address this possibility, we uncoupled the visual cues from the location of the cool spot as previously described ^17^. Specifically, the visual cues remained fixed while the cool spot location was changed. If flies in our assay were using non-visual cues for learning, they should progressively reduce their time to target even in this uncoupled condition. However, we find that in contrast to the coupled condition, *Cs* flies did not get faster at finding the cold-spot over time (Figure 2, M – O). Thus, flies in our assay use the distal visual cues to get progressively faster at locating the ‘cool spot’.

### Learning is sleep dependent

Sleep loss and extended waking result in cognitive deficits in a variety of tasks in animals from flies to humans ^3, 26, 27^. We therefore hypothesized that sleep deprivation would also impair spatial learning. To test this hypothesis, we sleep deprived *Cs* flies overnight using the Sleep Nullifying Apparatus ^20^, which deprived flies of >98% of their sleep (Figure 3A). As we expected, sleep deprivation impaired learning compared to age-matched controls (Figure 3B). Sleep deprivation does not alter simple visual behaviors such as object fixation and optomotor responses ^28^, suggesting that this was a learning defect rather than impaired visual acuity. To investigate whether sleep deprivation can independently alter heat avoidance, we placed flies in a chamber in which one half was heated at 36-37°C and the other half was maintained at 25°C. As seen in Figure 3C, sleep deprivation did not alter heat avoidance compared to untreated, age-match controls. Thus, the deficits in spatial learning following sleep deprivation are not due to alterations in sensory thresholds.

**Figure 3.**
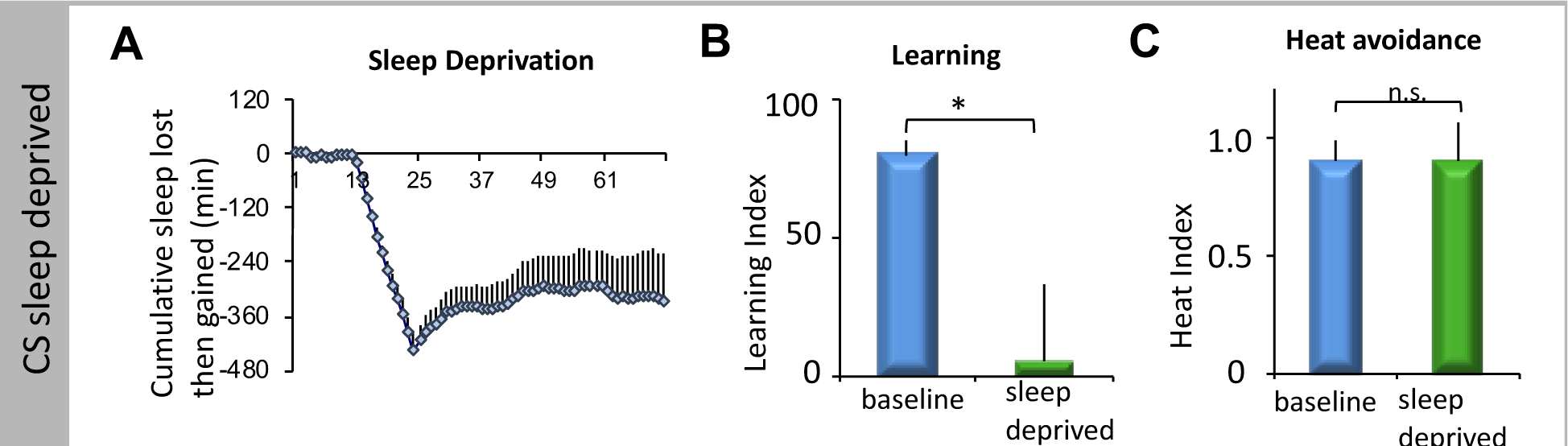
Sleep deprivation impairs spatial learning in *Cs* flies (A) Sleep deprived *Cs* flies lost 98% of their sleep and recovered ∼60% of their lost sleep during the subsequent 48hours in recovery (*n* = 30 flies, Repeated Measures ANOVA for time, F _[70,1470]_=12.97, *p* < 10^−15^). (B) Sleep deprived *Cs* flies (green) display impaired spatial leaning compared to controls (blue) (*n* = 7-10 flies / condition,* *p* < 0.01, t-test) (C) Sleep deprivation did not impair heat avoidance (*n* = 10-11 flies/ condition, n.s. *p* = 0.97, t-test).

### Enhancing sleep restores learning to *rutabaga* mutant flies

The adenyl cyclase *rutaba*ga (*rut*) was first identified as one of the canonical olfactory memory mutants in the fly ^29^. *rut* mutants have since been shown to be impaired in a number of different learning assays ^30-33^. Consequently, we hypothesized that *rut* mutants would also be impaired in spatial learning. As previously described, *rut*^*2080*^ mutants, which are in a *Cs* background, sleep the same as *Cs* controls ^19, 34^ (Figure 4A). Despite having similar sleep profiles, *rut*^*2080*^ mutants displayed severe behavioral impairment (Figure 4B). As with sleep deprivation, *rut*^*2*080^ are not impaired in optomotor responses ^35^ and exhibit normal heat avoidance (Figure 4C). Thus, *rut*^*2080*^ mutants display deficits in spatial learning. Enhancing sleep pharmacologically, by administering the GABA-A agonist Gaboxadol, can restore learning in *rut*^*2080*^ mutants when evaluated using a variety of learning assays including 1) Aversive Phototaxis Suppression assay, 2) courtship conditioning, and 3) place learning ^19, 34^ To determine whether enhanced sleep could also restore space learning to *rut*^*2080*^ mutants, we increased sleep in *rut*^*2080*^ mutants for two days by feeding them 0.1µg/mL Gaboxadol ^19, 36^. As shown previously, Gaboxadol-fed *rut*^*2080*^ males show a robust increase in total sleep time which is accompanied by a significant increase in the average duration of sleep bouts during the day (a measure of sleep consolidation) (Figure 4, D and E). Importantly, Gaboxadol-induced sleep significantly improved the learning index compared to age-matched, vehicle-fed siblings; Gaboxadol did not alter optomotor behavior (Figure 4, F and G). Taken together, these results support and extend our previous observations that enhancing sleep can reverse the learning impairments in the classic memory mutant *rut* ^19^.

**Figure 4.**
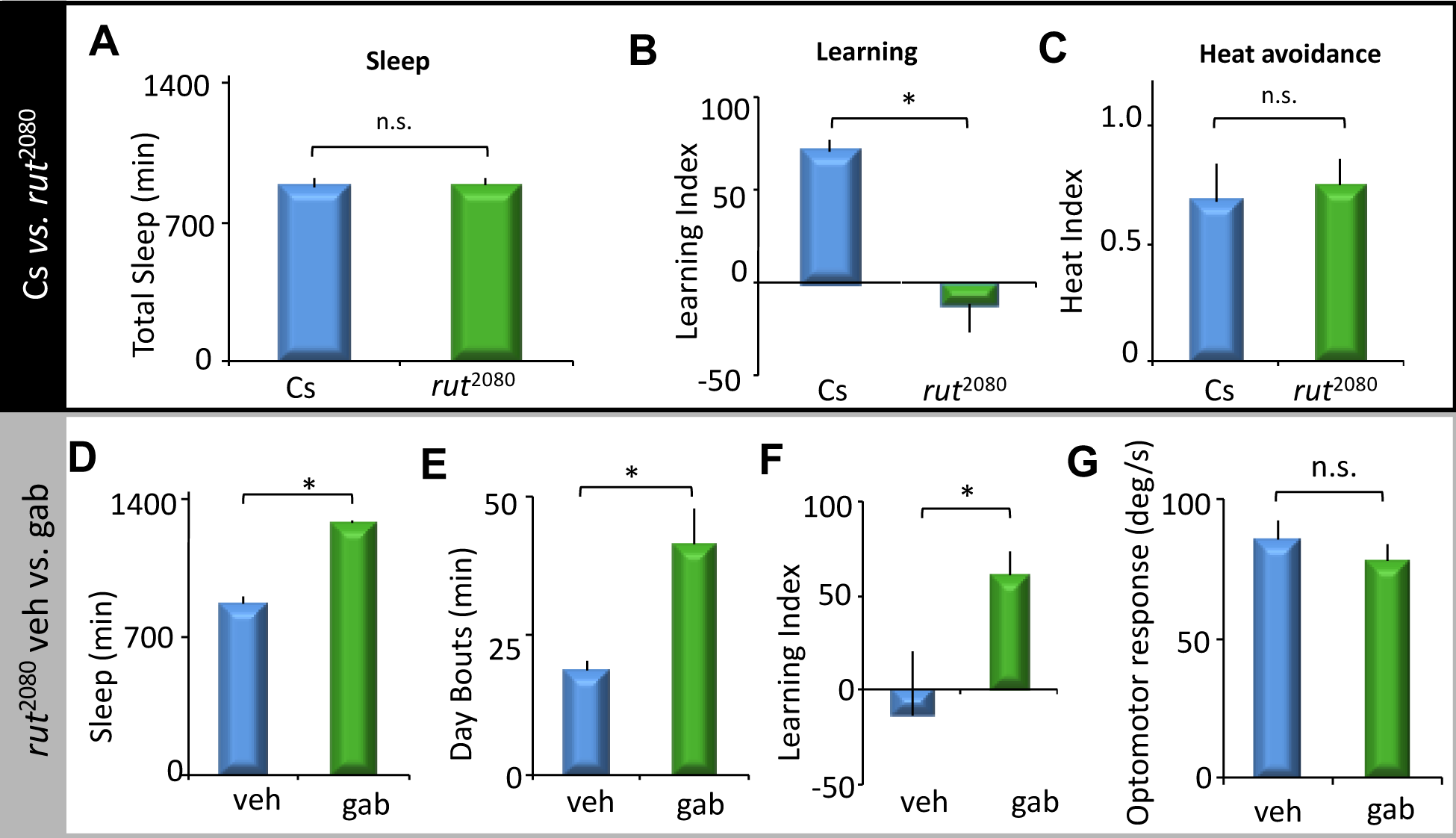
*rut* dependent impairments in spatial learning are reversed by enhancing sleep (A) Sleep profiles of *rut*^2080^ mutants was not different compared to *Cs* controls (*n* = 18-22 flies / genotype, n.s. *p* =0.49, t-test). (B) Spatial learning is impaired in *rut*^2080^ mutants compared to *Cs* controls (*n* = 9-10 flies / genotype,* *p* < 0.001, t-test). (C) *rut*^2080^ males are not impaired in heat avoidance (*n* = 10 flies / genotype, n.s. *p* = 0.27). (D) Gaboxadol increases total sleep in *rut*^2080^ flies compared to vehicle-fed siblings (*n* = 18-20 flies/condition, * *p* < 10-^10^, t-test). (E) Gaboxadol –fed *rut*^*2080*^ flies display increased average daytime sleep bout duration compared to vehicle fed siblings (* *p* < 10^−4^, t-test). (F) Gaboxadol restored spatial learning to *rut*^*2080*^ flies compared to controls (*n*= 9-10 flies / condition,* *p* < 0.01, t-test). (G) Gaboxadol did not impair the optomotor response of *rut*^*2080*^ flies (*n*= 19 flies / condition, n.s. *p* = 0.43, t-test).

### Age-dependent learning impairments are reversed by enhancing sleep

Age-dependent decline in cognitive performance has been observed in flies and humans ^37-40^. In flies, some plasticity deficits are observed as early as 18-20 days of age ^37, 40^. We therefore hypothesized that flies would exhibit age-dependent degradation in spatial learning as well. We started by examining sleep in 21-24 day old flies. As previously described, older flies had less total sleep and shorter average sleep bout duration than 4-5 day old flies (Figure 5 A-C) ^22^. Importantly, waking activity of old flies was not altered indicating that locomotor activity was not impaired (Figure 5D). Interestingly, 21-24 day old flies displayed impairments in spatial learning compared to 4-5 day old flies (Figure 5E). As above, the changes in performance were not associated with impairments in heat avoidance or visual acuity, indicating that the learning defects we observed in old flies were not a consequence of defective sensory processing (Figure 5, F and G). Thus, like amnesiac-dependent memory and Social enrichment, 20 day old flies also show deficits in space learning ^37, 40^.

**Figure 5.**
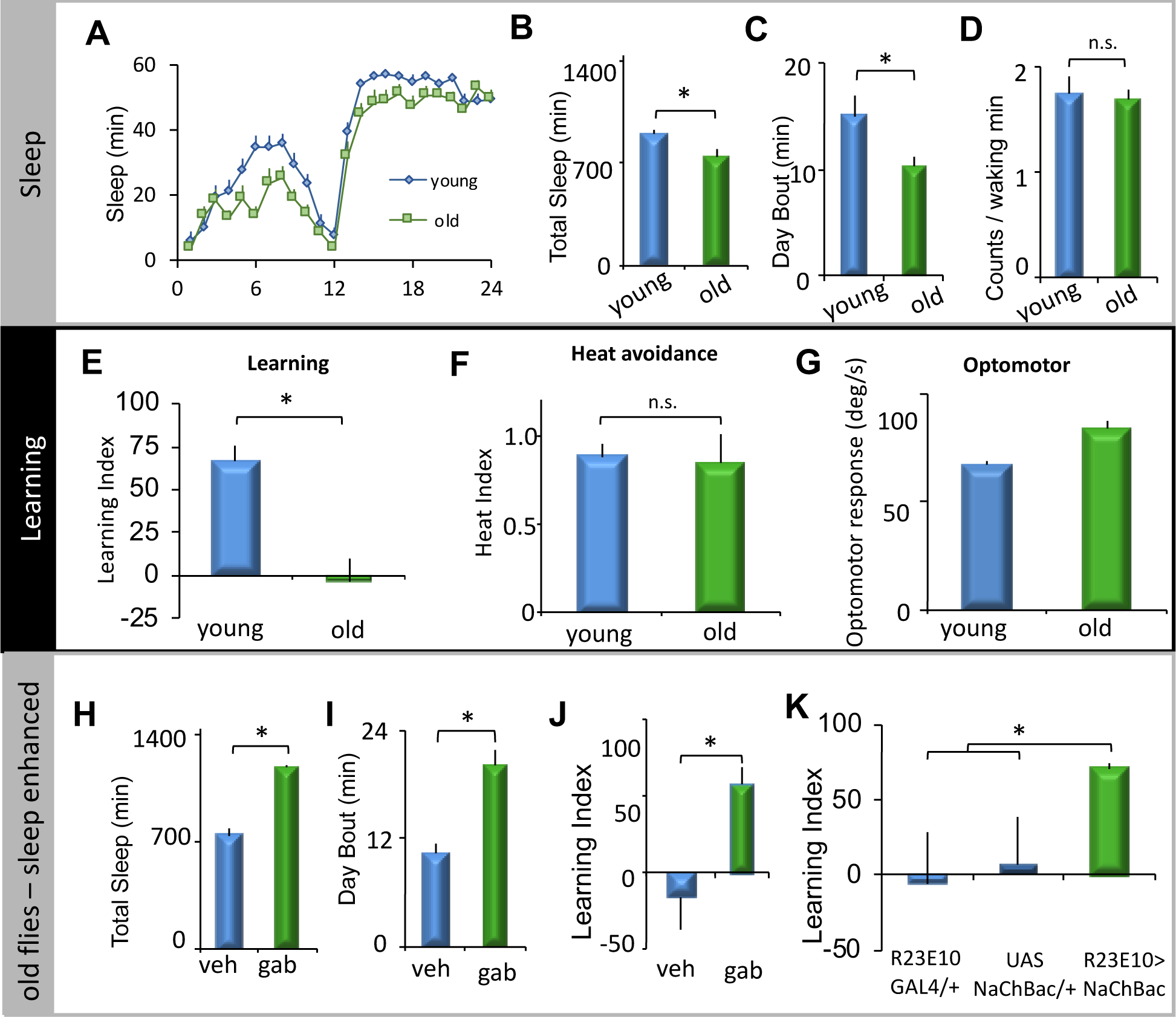
Age dependent declines in spatial learning can be reversed by enhancing sleep. (A) Sleep, in minutes per hour, was reduced in 21-24 day old flies (green) compared to 5 day old controls (blue) (*n*= 20 – 27 flies / group, repeated measures ANOVA age X time; F_[23,966]=_5.49, *p* < 0.001). (B) Total sleep was reduced in 21-24 day old flies compared to 5 day old flies;* *p* < 0.01, t-test. (C) Aging reduced average daytime sleep bout duration; *p* < 0.05, t-test. (D) Waking activity was not impaired in 21-24 day old flies (n.s. *p* = 0.83). (E) Spatial Learning was impaired in 21-24 day old flies compared to 5 day old flies (*n* = 10-14 flies / group, * *p* < 10^−4^). (F) Age did not disrupt heat avoidance (F, *n* = 10 flies / condition, n.s. *p* = 0.49). (G) Age did not disrupt optomotor responses (G, *n* = 51-62 flies / condition, * *p* < 10^−5^). (H) Gaboxadol (Gab) increased sleep in 21-24 day old flies (H, *n* = 20 flies / group, * *p* < 10^−10^). (I) Gaboxadol increased average daytime sleep bout duration in 21-25 day old flies (green) compared age-matched controls (blue) (* *p* < 10^−4^, -t-test). (J) Spatial learning was restored to Gaboxadol-fed 21-24 day old flies (green)compared to age-matched vehicle fed controls (*n* = 9-10 flies / condition, * *p* < 10^−4^, t-test). (K) Spatial learning was significantly higher in 21-24 day old R23E10-GAL4/+>UAS-NaChBac/+ flies compared to age matched R23E10-GAL4/+ and UAS-NaChBac/+parental controls (*n* = 8-10 flies / genotype, One way ANOVA for genotype F_[2,49]_=4.59, *p* < 0.05, * *p* < 0.01, modified Bonferroni test)

Given that older flies show sleep deficits (Figure 5, A-C) and previous reports have found that genetically enhanced sleep can restore plasticity to older flies, we hypothesized that enhancing sleep might also restore spatial learning to 20-24 day old flies ^40^. We tested this hypothesis by enhancing sleep in old flies with two different methods – pharmacologically, by feeding old flies Gaboxadol, and genetically, by activating the fan shaped body (a known sleep center)^41^. Gaboxadol robustly increased sleep amount and consolidation in old flies (Figure 5, H and I). Crucially, Gaboxadol-enhanced sleep restored memory to 20-day old flies compared to vehicle-fed, age-matched siblings (Figure 5J). Gaboxadol did not alter heat avoidance in 20-day old flies (see below) indicating that the improvements were not due to changes in sensory thresholds. To confirm these results, sleep was increased by expressing the bacterial sodium channel NaChBac under the control of the R23E10-GAL4 driver. Consistent with previous results, activating the Fan Shaped body increased sleep (data not shown). Importantly, 20-day old *R23E10-GAL4/+>UAS-NaChBac/+* flies displayed significantly higher learning scores than both parental controls (*R23E10-GAL4/+* and *UAS-NaChBac/+*) (Figure 5K). Thus, inducing sleep with two independent methods reverses the age dependent cognitive deficits we see using spatial learning. These results support previous suggestions that sleep can be used as a therapeutic to reverse age dependent cognitive deficits ^42, 43^.

### Spatial learning requires dopamine signaling

The neuromodulator dopamine plays key roles in facilitating synaptic mechanisms that support learning and memory in flies and mammals ^20, 44-47^. However, the potential role of dopaminergic signaling in spatial learning in flies has not yet been investigated. Thus, we evaluated spatial learning while using both pharmacology and genetics to modulate dopamine. We disrupted dopamine pharmacologically by feeding flies the dopamine synthesis inhibitor 3-Iodo L Tyrosone (3IY) ^48^. As seen in Figure 6 A-C, feeding flies 3IY increased both total sleep time, and sleep consolidation during the day compared to age-matched vehicle-fed siblings; no changes in waking activity were observed (Figure 6D). These data highlight the wake-promoting effects of dopamine^50^. Importantly, feeding flies 3IY impaired learning (Figure 6, E – G), suggesting that spatial learning requires dopaminergic signaling.

**Figure 6.**
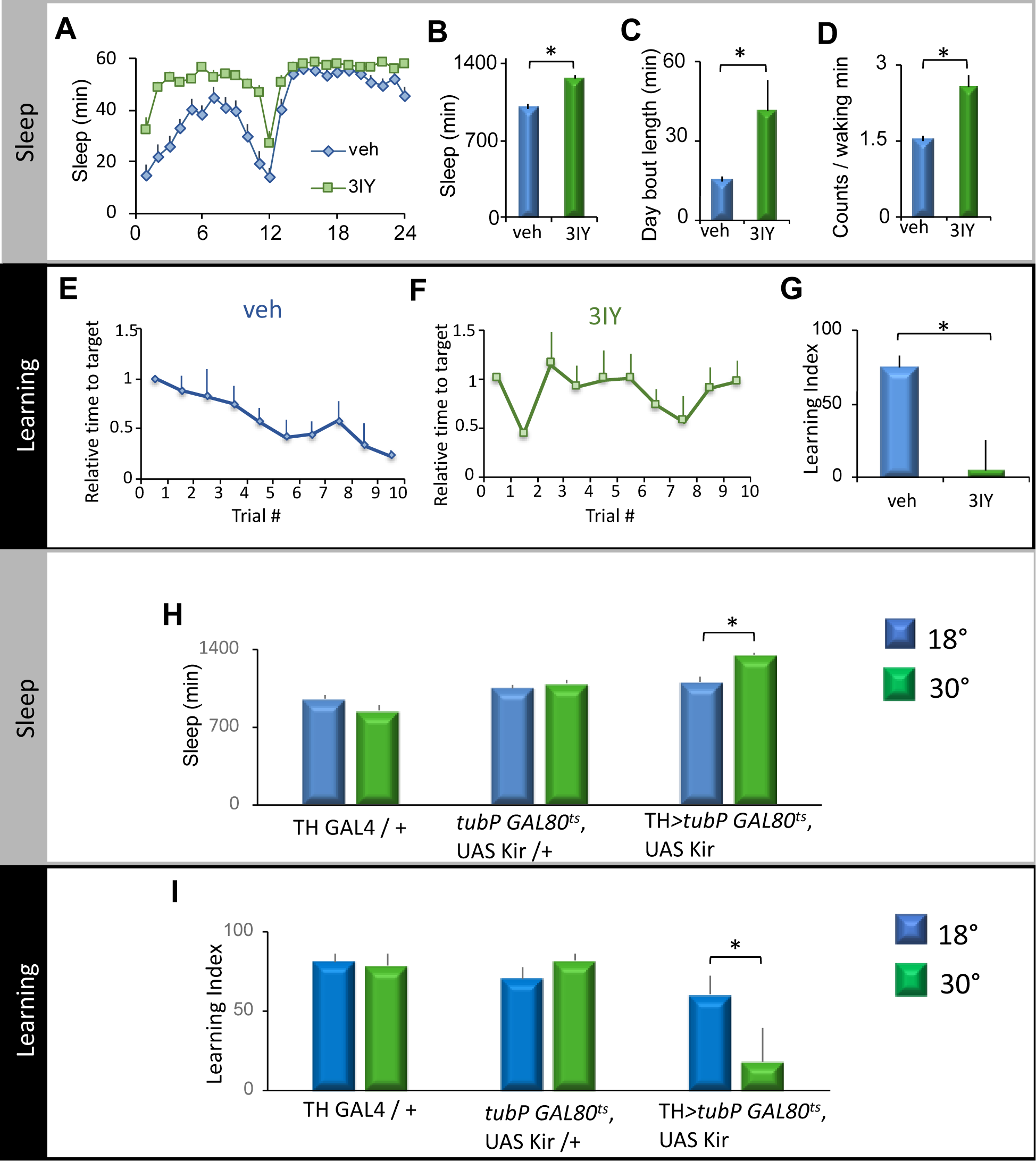
Reducing dopamine signaling impairs learning. (A) 3-Iodo-L-Tyrosine (3IY)-fed flies display an increase in sleep compared to vehicle fed controls (Sleep in minutes per hour, *n* = 20-21 flies / group, Two way repeated measures ANOVA for Drug X Time F _[23,782]_=5.49,*p* < 10^−6^). (B) 3IY fed flies display more total sleep than age-matched vehicle fed siblings (* *p* < 10^−6^, t-test). (C) 3IY increased average daytime sleep bout duration (* *p* < 0.001, t-test). (D) 3IY did not impair waking activity compared to vehicle fed controls (* *p* < 0.01, t-test). (E) 3IY-fed *Cs* flies were impaired in spatial learning. (F) In contrast to 3IY –fed flies, vehicle fed controls displayed spatial learning (*n* = 8 flies/group, Two way ANOVA Drug X Trial, F _[9,126]_=2.33, *p* < 0.05). (G) Learning index of 3IY fed flies was greatly reduced compared to vehicle-fed controls (* *p* < 0.01, t-test). (H) *TH-GAL4/+>GAL80*^*ts*^, *UAS Kir/+* flies displayed an increase in sleep at 30°C compared to siblings maintained at 18°C; sleep in *TH GAL4* / *+* or the *tubP GAL80*^*ts*^, *UAS Kir* /*+* parental controls was similar at both 18°C and 30°C (*n* = 20-30 flies / group, Two way ANOVA for genotype X temperature, F_[2,131]_=7.28, *p* < 0.01; * *p* < 0.001, modified Bonferroni test). (I) Spatial learning is impaired in *TH>GAL80*^*ts*^, *UAS Kir* flies at 30°C compared to siblings maintained at 18°C; temperature did not impact spatial learning in either *TH GAL4* / + or the *tubP GAL80*^ts^, *UAS Kir* /+ parental controls (*n*= 8-12 flies / group, Two way ANOVA for genotype X temperature, F _[2,116]_=4.96,*p* < 0.01, * *p* < 0.05, modified Bonferroni test).

To disrupt dopamine genetically, we inhibited most dopaminergic neurons by expressing the inwardly rectifying potassium channel KCNJ2 (UAS-Kir2.1) using Tyrosine Hydroxylase GAL4 (TH-GAL4). To confine the inhibition of dopaminergic neurons to the adult stage, and obtain better temporal control of inhibition, we used the TARGET system. The TARGET system uses a temperature sensitive GAL4-suppressor, *GAL80*^ts^. *GAL80* is inactivated, thereby relieving the suppression of GAL4, and allowing the expression of UAS-Kir2.1 only at 30°C ^49^. In support of this interpretation of the 3IY results, at 30°C, *TH GAL4* > *GAL80*^ts^; *UAS Kir 2*.*1* flies displayed both increased sleep (Figure 6H), and impaired spatial learning (Figure 6I) compared to siblings maintained at 18°C; *TH-GAL4/+* and *tubP GAL80*^ts^, *UAS Kir /+* parental lines displayed normal sleep and memory at 18°C and 30°C. Thus, reducing dopamine levels with two different methods impairs spatial learning indicating that dopaminergic signaling is required for learning in this assay. Dopamine deficient flies have normal optomotor responses, visual fixation, and electroretinograms indicating that the spatial learning impairments were not a consequence of aberrant sensory processing ^51^.

### Enhancing dopamine signaling reverses age dependent cognitive impairments

Dopamine levels are known to decrease with age in flies, even as dopaminergic neurons appear to be anatomically unaffected ^52, 53^. Further, we have previously shown that enhancing dopamine signaling in 20-day old flies restores structural age-dependent deficits in behavioral plasticity ^40^. Therefore, we hypothesized that enhancing dopamine signaling would reverse the age dependent spatial learning impairments observed above. Dopamine was increased by feeding flies the dopamine precursor Levodopa (L-Dopa)^54, 55^. As seen in Figure 7A, feeding 20-day old flies L-DOPA disrupted nighttime sleep as previously reported ^56, 57^. Importantly, spatial learning was restored in 20-day old, L-Dopa fed flies compared to their age-matched vehicle-fed siblings (Figure 7B). Further, feeding L-Dopa or Gaboxadol did not alter heat avoidance (Figure 7C).

**Figure 7.**
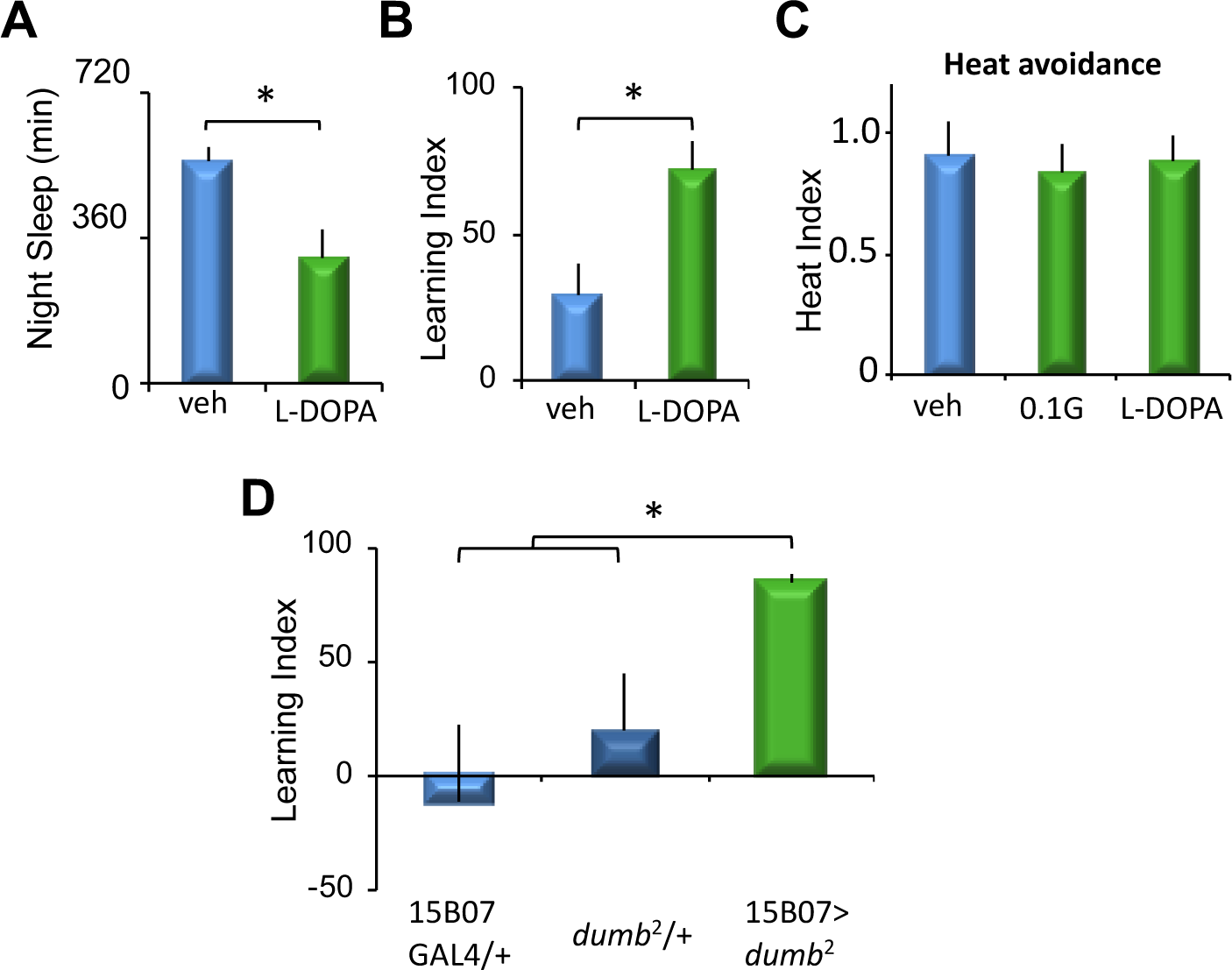
Enhancing dopamine signaling ameliorates age related cognitive decline. (A) Night-time sleep was reduced in Levodopa (L-Dopa, green) fed 21-24 day old *Cs* flies compared to age-matched vehicle-fed (blue) controls (*n =* 10-12 flies / group, * *p <* 0.01, t-test). (B) Spatial learning was elevated in old L-Dopa fed *Cs* flies (green) compared to age-matched controls (blue) (*n* = 9-10 flies/group, * *p* < 0.01). (C) Heat avoidance was not changed in old flies fed L-Dopa or Gaboxadol (green) compared to age-matched controls (*n* = 10 flies / condition, n.s. p > 0.25, modified Bonferroni test). (D) Spatial learning was elevated in old *R15B07*>*dumb*^2^ flies compared to age-matched *R15B07 GAL4*/ + and *dumb*^2^ / + parental controls (*n* = 8 flies / genotype, One way ANOVA for genotype F_[2,45]_=5.82, *p* < 0.01; * *p* < 0.01, planned comparisons, modified Bonferroni test).

Spatial learning in flies is known to require the function of R1 ellipsoid body (EB) ring neurons ^17^. Further, dopamine receptors are known to be expressed in the EB ^58, 59^. Combined with our results above showing that we could restore learning to aged flies by elevating dopamine levels, we hypothesized that age-dependent learning impairments could be reversed by elevating dopamine signaling in the EB. To test this hypothesis, we expressed the *Drosophila Dopamine D1 Receptor* (*Dop1R1*) in the EB using R15B07-GAL4. The *Dop1R1* mutant, *dumb*^*2*^, contains a piggyBac inserted into the first intron of the *Dop1R1* gene that contains a UAS that can be used to induce a functional *Dop1R1* receptor ^60^. As seen in Figure 7D, disruptions in spatial learning are reversed in 20-day old R15B07-GAL4/+>*dumb*^2^/+ flies compared to age-matched parental controls (*15B07-GAL4/+* and *dumb2/+*). Thus, increasing dopaminergic signaling through the Dop1R1, specifically in the R1 ellipsoid body ring neurons rescues age-dependent cognitive decline in spatial learning.

## DISCUSSION

We find that sleep plays an important role in supporting spatial learning in flies. Sleep deprivation impaired learning; conversely, enhancing sleep reversed learning impairments associated with *rut* mutants and aging. These data build on previous results that suggested a surprising restorative depth to the relationship between sleep and plasticity ^19, 34^, and extend them to a novel spatial learning task. As discussed below, our results are consistent with findings in rodent spatial learning and human episodic memory research, reinforcing the parallels between spatial learning in animal models and human declarative memories ^15, 16^.

The modified spatial learning assay studied here was adapted from previous designs ^17, 18^. In the published protocols, 8-10 replicates using 15 flies per replicate are tested using a between subject design. In contrast, we chose to study individual flies to allow us to track learning in each fly using a within subject design. Testing groups of 15-100 flies/replicate is standard in *Drosophila* learning and memory studies and provides many advantages ^29, 61^. However, testing individual flies in a within subject design provides additional opportunities. For example, by studying the learning behavior of individual flies, we can isolate and study flies that display a range of phenotypes (e.g. different rates of learning). We have previously used this approach to identify genes that convey resilience or vulnerability to sleep loss.^62^. Further, evaluating individual flies substantially reduces the computational power needed to evaluate details of behavior (path length etc.). Moreover, testing individual flies also reduces the amount of time required to generate the necessary flies to complete a given experiment and can expedite discovery experiments. Importantly, examining spatial learning in 8-10 flies/genotype produces statistically robust datasets. Indeed, significant learning was observed in two independent replicates of two different wild-type strains – *Cs* and 2U, thus validating our design for spatial learning.

### Spatial Learning is Sleep Dependent

Depriving flies of sleep overnight impaired spatial learning. These results are consistent with experiments in rodents and humans. In rodents, sleep deprivation impaired encoding of hippocampus-dependent spatial memory as assessed with the Morris Water Maze, while sleep loss had minimal effects on hippocampus independent non-spatial tasks ^63, 64^. These experiments in rodents are corroborated by studies in humans that found that human spatial memory was dependent on sleep ^65^. Further, sleep deprivation in humans also impaired learning in declarative memory tasks which are known to require hippocampus function ^12, 13^.

### Enhancing sleep restores spatial learning to *rut* mutants

The *rutabaga* (*rut*) mutant was first isolated as one of the canonical fly learning and memory mutants using olfactory conditioning ^29^. *rut* mutants have since been shown to be impaired in a number of different learning and memory assays, and have been used to validate new assays ^20, 29, 31-33^. We therefore evaluated spatial learning in *rut* mutants. Indeed, we find that *rut* mutants don’t exhibit any sleep defects, but nonetheless, are severely impaired in spatial learning. Moreover, enhancing sleep pharmacologically by feeding *rut* mutant flies Gaboxadol for two days restored spatial learning. These results are consistent with previous work showing that Gaboxadol enhanced sleep restored learning to *rut* mutants in other learning assays: Aversive Phototaxis Suppression, courtship conditioning, and place learning ^19, 34^.

Importantly, *rut* mutants do not display sleep defects during baseline. As a consequence, it is unlikely that Gaboxadol-induced sleep is simply ameliorating preexisting sleep deficiencies. Rather, the enhanced sleep induced by Gaboxadol is likely to exert its effects on neuronal plasticity in memory circuits. For olfactory conditioning, *rut* has been proposed to function as a coincidence detector in mushroom body Kenyon cells, detecting coincident input of the conditioned stimulus (odor) and the unconditioned stimulus (electric shock)^66^. However, *rut* is widely expressed, and likely functions as a signaling molecule in multiple cellular processes to influence different aspects of neural plasticity ^19, 35, 67-72^. Indeed, it should be noted that the brain processes odors using sparse coding and that, during olfactory conditioning, the electric shock is very precisely timed with brief puffs of odors to induce a lasting association ^29, 73^. In contrast, flies evaluated using many operant learning assays, such as courtship conditioning, place learning and Aversive Phototaxic Suppression, experience a more continuous exposure to the aversive stimulus (quinine, mate-rejection and heat) ^30, 74-76^. In any event, the precise role of *rut* in spatial learning, its site of action, and the mechanism by which enhanced sleep restores learning to *rut* mutants remain unknown and are the subject of ongoing study.

### Enhancing sleep reverses age dependent cognitive decline

Age-related memory impairments are observed in humans, and appear to disproportionately affect hippocampus dependent episodic memories and spatial memory ^77-79^. Further, aging is also accompanied by sleep deficits and defects in sleep dependent memory consolidation ^5, 80^. Enhancing sleep in older adults was also able to ameliorate age-related impairments ^42, 43^.

Flies too, have been shown to exhibit age-dependent cognitive decline ^37, 38, 40, 81^. In some cases, plasticity deficits have been observed at 18-20 days of age ^37, 40^. Further enhancing sleep in aged flies by feeding Gaboxadol reversed age-dependent defects in social-enrichment induced plasticity ^40^. Consistent with these results, we found that 20-24 day old flies were impaired in spatial learning and that enhancing sleep could reverse these impairments. Interestingly, the age-dependent spatial learning impairments we observed appear to be more severe than those observed with olfactory conditioning ^37^. These data parallel studies in rodents that found that aging impaired hippocampus dependent spatial learning but did not appear to affect hippocampus independent non-spatial learning ^82-84^.

### Dopamine signaling is required for spatial learning

We find that inhibiting dopaminergic signaling with two different methods increased sleep and impaired spatial learning. In rodents, dopamine secreted from locus coeruleus to the hippocampus also plays a critical role in mediating spatial learning ^85-87^. Our data, thus support a conserved role for dopamine in spatial learning, consistent with its role as a key facilitator of synaptic plastic changes that support learning and memory ^44-46^.

It is worth noting that both methods of inhibiting dopaminergic signaling increased sleep while impairing learning. We have argued previously that a thorough characterization of sleep should not rely exclusively on examining sleep metrics only. Healthy sleep promotes a number of positive non-sleep variables such as memory, plasticity, metabolism, immune function, etc.

Determining whether a change in sleep induced by a genetic manipulation impacts these other variables is essential for understanding whether sleep has been positively or negatively impacted ^26, 62^. Our data clearly indicate that the increased sleep associated with impaired dopaminergic signaling is associated with impairments in spatial learning. Given that Gaboxadol-induced sleep restores spatial learning to *rut*^*2080*^ mutants and 20-21 day old flies, we hypothesize that disrupting dopamine signaling disrupts sleep efficiency. Indeed, while *dumb*^2^ flies sleep more, they are also more arousable at night, suggesting they are not sleeping as deeply ^88^. That is, the flies would need to sleep more to compensate for this ineffective sleep.

Another non-exclusive hypothesis is that different subsets of dopaminergic neurons support arousal and spatial learning. The fly arousal promoting dopaminergic neurons are known to project to the fan shaped body and the mushroom body ^89-91^. The dopaminergic neurons that support spatial learning are not yet known. Our spatial learning assay requires the function of EB ring neurons ^17^. Dopamine receptors are known to be expressed in the EB ^58, 59^. Further, dopaminergic neurons have been described that project into the EB ^59, 92^ from the PPM3 cluster. These PPM3 EB projecting dopaminergic neurons are great candidates for mediating spatial learning. Thus, the effects of dopaminergic inhibition on sleep and learning could map to different subsets of dopaminergic neurons. Further experiments are needed to distinguish between these possibilities. Nonetheless, the apparent discord between increasing sleep and impairing learning serves to highlight the importance of functional evaluation of sleep outcomes when describing manipulations that affect sleep time ^26^.

### Enhancing dopaminergic signaling reverses age dependent impairment

Complimentary to the experiments above where we inhibited dopaminergic signaling, we find that increasing dopaminergic signaling restored spatial learning to aged flies. These results are consistent with previous work in flies and mammals. Dopamine levels decline with age in flies and humans ^52, 93, 94^. Further, enhancing dopaminergic signaling reversed aspects of age-dependent cognitive decline. In humans, elevating dopaminergic signaling ameliorated age-dependent declines in episodic memory ^95^. Similarly, increasing dopaminergic signaling restored spatial learning in rodents ^84^, and reversed age dependent defects in social-enrichment induced plasticity in flies ^40^. How is enhancing dopaminergic signaling able to restore learning to aged brains? Overexpressing the Dop1R1 dopamine receptor in EB ring neurons restored learning to aged flies. This result suggests the possibility that aging disrupts dopaminergic signaling to the EB (at the level of dopamine release and/or receptivity). This possibility will be addressed in future work.

## CONCLUSIONS

Collectively our results demonstrate a critical role for sleep in supporting spatial learning in flies. Sleep deprivation impairs space learning. Conversely, enhancing sleep restores learning to impaired brains. Our data are consistent with work on spatial learning in rodents and spatial and episodic memories in humans, indicating that the phenomena we report are conserved.

Interestingly, spatial learning in mammals is dependent on the hippocampus, and is closely associated with the phenomenon of replay. When animals are trained to run along a linear track, their trajectories are represented by a sequence of activation of hippocampal place cells ^96^. These sequences are replayed in a time-compressed fashion during sleep ^97-99^, in a complex dialog between the hippocampus and the cortex ^100-103^, to consolidate the memory of the experience ^104, 105^. Further, these place cell sequences can be reactivated by cueing in sleep ^103, 106-108^. Although replay like phenomena have not been observed in flies, cued reactivation during sleep improved recall in bees ^109^, and reactivation during sleep of dopaminergic neurons involved in memory acquisition was shown to facilitate consolidation of courtship memory in flies ^110^, suggesting that such replay-like processes might be detected in *Drosophila* too.

## ACKNOWLEDGEMENTS

We thank Lijuan Cao for technical assistance, Michael Reiser for helpful advice and technical input, and Matt Thimgan and Stephane Dissel for a critical reading of the manuscript.

